# Soluble SorCS1 binds the insulin receptor to enhance insulin sensitivity

**DOI:** 10.1101/2020.11.10.376178

**Authors:** Mads Kjolby, Peter Breining, Karen Marie Pedersen, Niels Wellner, Karen Marie Juul Sørensen, Anette Marianne Prior Gjesing, Dovile Januliene, Dorthe Maria Vittrup, Arulmani Manavalan, Maarten Rotman, Gangadaar Thotakura, Guido Hermey, Peder Madsen, Christian B. Vaegter, Søren Thirup, Olof Asplund, Rashmi B. Prasad, Edwin T. Parlevliet, Patrick C.N. Rensen, Andreas Nygaard Madsen, Birgitte Holst, Olav M. Andersen, Arne Möller, Torben Hansen, Anders Nykjaer

**Author notes:** Corresponding authors Anders Nykjaer, PROMEMO and DANDRITE, Department of Biomedicine, Hoegh-Guldbergsgade 10, Aarhus University, DK-8000 Aarhus C, Denmark, Mads Kjolby, DANDRITE, Department of Biomedicine, Hoegh-Guldbergsgade 10, Aarhus University, DK-8000 Aarhus C, Denmark. Shared first authors (Mads Kjolby, Peter Breining).

## Abstract

Type 2 diabetes has reached endemic proportions and is a substantial burden for the affected patients and the society. Along with lifestyle factors, a number of genetic loci predisposing to type 2 diabetes have been identified, including *SORCS1* that encodes the transmembrane receptor SorCS1. The ectodomain of SorCS1 (sol-SorCS1) is shed from plasma membranes but the biological function of this fragment is unknown. Here we show that sol-SorCS1 acts as a high-affinity binding partner for the insulin receptor to stabilize the receptor and increase insulin affinity, protein kinase B activation, and glucose uptake in myocytes. Sol-SorCS1 is liberated from adipocytes, and in diabetic patients the plasma concentration positively correlates with body mass index, but inversely with plasma glucose. In mouse models of insulin resistance, exogenous sol-SorCS1 restored insulin sensitivity. We conclude that sol-SorCS1 increases peripheral insulin sensitivity and propose sol-SorCS1 as a novel insulin sensitizing adipokine and potential antidiabetic agent.

---

The mechanisms controlling insulin sensitivity and glucose metabolism are fundamental to all vertebrates. Insulin resistance is an early hallmark for the development of type 2 diabetes^1,2^ as insulin sensitivity and peripheral tissue glucose uptake is essential to acutely buffer plasma glucose. The downstream signaling pathways of the insulin receptor (IR) in skeletal muscle are well described^3^. Among the second messengers, the most prominent pathway regulating the insertion of GLUT4-containing vesicles is the protein kinase B (AKT) signaling cascade^3^. However, whether this pathway can be dynamically regulated through interaction with co-receptors or other membrane proteins is unknown^4-6^. The existence of a molecular thermostat to adapt insulin sensitivity to the metabolic state would transform our conceptual understanding on how key metabolic processes can be controlled.

SorCS1 is a member of the Vps10p-domain family of sorting receptors that also comprises sortilin, SorLA, SorCS2, and SorCS3^7-10^. Several studies have identified *SORCS1* as a susceptibility locus for type 2 diabetes mellitus that is conserved between humans^11-13^, rats^14^ and mice^15^. Recently, intracellular SorCS1 was shown to facilitate the biogenesis of insulin containing secretory granules, but a function outside the β-cells was not studied^16^. We therefore performed RNA sequencing on human samples focusing on major peripheral tissues involved in glucose homeostasis (**Fig 1,a**). Notably, we found that SorCS1 expression was highest in adipose tissue surpassing that of the pancreatic islets whereas expression in the liver was hardly detected. We previously showed that every hour up to 95% of SorCS1 (sol-SorCS1) is liberated from the plasma membrane by ADAM17/TACE mediated cleavage but the biological relevance of this observation is unknown^17,18^. Given that human and murine adipose tissue express ADAM17/TACE^17^, we examined, whether body mass index (BMI) associates with the plasma concentration of sol-SorCS1. To this end we assessed the correlation between BMI and the concentration of sol-SorCS1 in plasma in a cohort of diabetic patients and healthy controls. We found that sol-SorCS1 positively correlated with BMI and most prominently did so in the diabetic cohort (**Fig 1,b**). Accordingly, sol-SorCS1 could be liberated from human adipocytes as it was clearly detected in the conditioned medium from human subcutaneous adipose tissue explants (**Fig. 1,c**). By immunoprecipitation (IP) we also detected SorCS1 expression in lysates of visceral fat from mice and differentiated murine 3T3-L1 adipocytes, respectively (**Fig. 1,d-e**). Notably, the conditioned medium of the 3T3-L1 cultures was enriched in the soluble form of SorCS1 that migrated slightly faster than the full-length receptor present in the corresponding lysate. Although SorCS1 protein was found in the myocyte-derived murine cell line C2C12 (**Extended data 1,a**), we failed to detect the soluble receptor in the conditioned media (**Extended data 1,b**). We next examined whether sol-SorCS1 levels is associated with plasma glucose. Remarkably, among the diabetic patients sol-SorCS1 inversely correlated with plasma glucose (**β**-value -2.6E-3, P<0.001) (**Fig 1,f**), suggesting that the soluble receptor may impact glucose metabolism.

**Figure 1:**
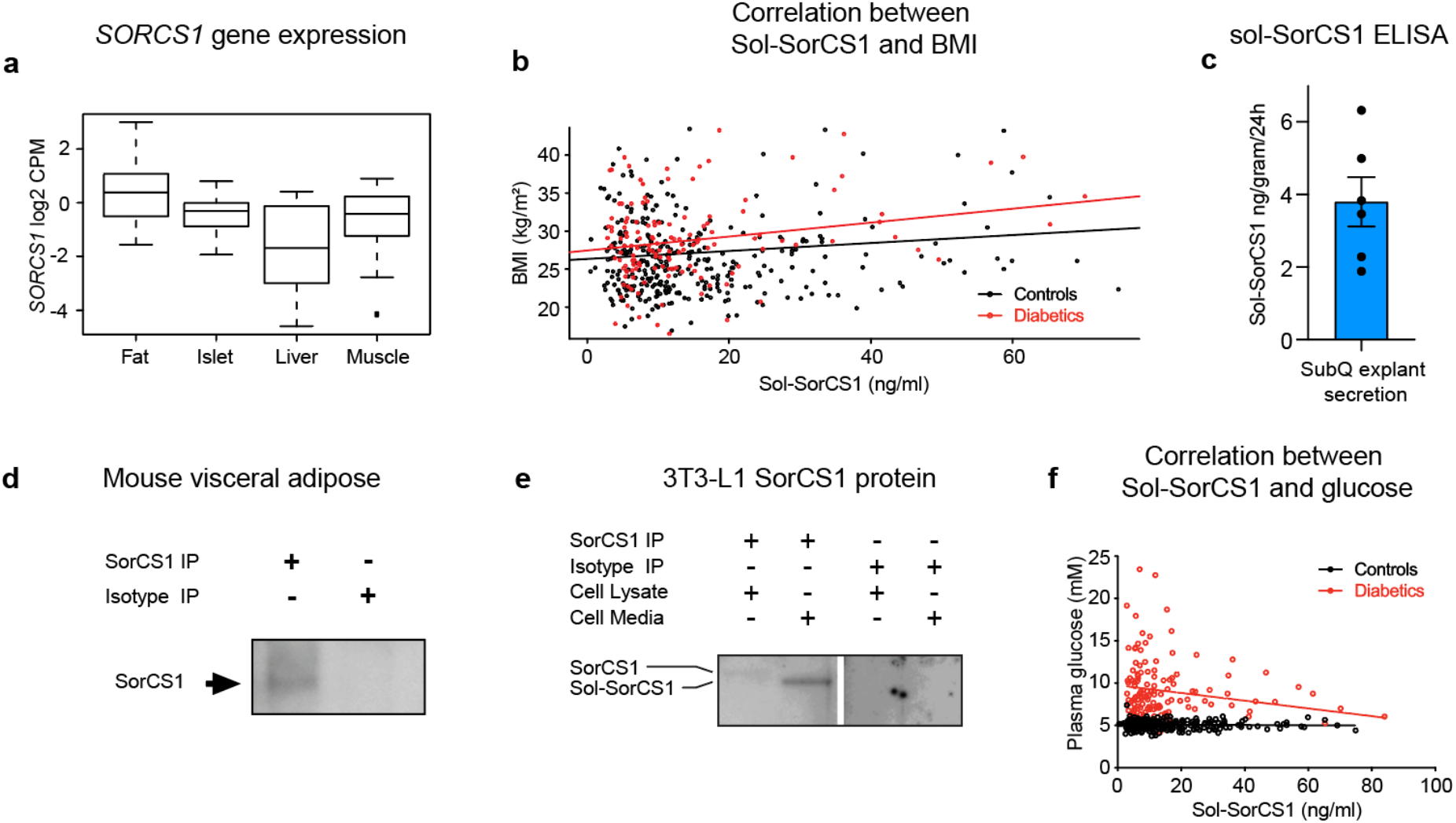
SorCS1 is expressed in, and secreted from, adipose tissue **a**, mRNA expression of *SORCS1* in human tissues (n=12/group). **b**, in a group of healthy controls and diabetes patients (n=328 and n=124, resp.) plasma levels of sol-SorCS1 positively correlate with BMI (**β**-value 0.011, P=0.007 and **β**-value 0.058, P<0.05, respectively). **c**, Sol-SorCS1 ELISA of media from 24h incubation of human subcutaneous adipose tissue (SubQ) explants (3.80±0.68 ng/gram/24h). **d**, protein immunoblot of murine SorCS1 in adipose tissue. **e**, full-length SorCS1 and its soluble shed form in 3T3-L1 adipocytes. **f**, in the human cohort sol-SorCS1 negatively correlates with fasting plasma glucose (**β**-value -2.6E-3, P<0.001).

We hypothesized that sol-SorCS1 could serve as an adipose tissue derived peptide hormone to compensate for an increase in BMI by lowering plasma glucose levels and preventing hyperglycemia. To test this hypothesis, we sought to assess a potential glycemic regulatory function of the soluble receptor. We therefore injected an adenovirus encoding human sol-SorCS1 (AV-sol-SorCS1) or AV-LacZ (encoding β-galactosidase), as a negative control, into the tail vein of diet-induced obese (39.5±1.2g) hyperglycemic C57BL/6 mice (DIO mice). Eleven days after viral transduction, overexpression of the soluble receptor substantially mitigated the diabetic phenotype by lowering both the elevated plasma glucose and insulin levels corresponding to a 24% decrease in plasma glucose and a 22% reduction in insulin levels, respectively (**Fig 2,a-b**). To test the antidiabetic efficacy of sol-SorCS1 in another and more severe diabetic model, we turned to the leptin receptor knockout mouse model (*db/db*) that rapidly develops progressive obesity and type 2 diabetes. One week after injection with AV-LacZ, plasma glucose was increased by 55% and insulin by 65% due to the gradual worsening of the diabetic phenotype (**Fig. 2,c-d**). Remarkably, following AV-sol-SorCS1 treatment plasma glucose remained unchanged and this despite a decrease in plasma insulin levels, indicating increased insulin sensitivity. In the intraperitoneal glucose tolerance test (IPGTT), AV-sol-SorCS1 mice responded markedly stronger with lower plasma glucose levels at all timepoints corresponding to a reduction in the area under the curve (AUC) by 79.5% (**Fig. 2,e-f)**. The treated animals also exhibited superior ability in mobilizing insulin compared to the control group, revealing a spare capacity of insulin production that the control animals did not possess (**Fig. 2g)**. Notably, regardless the level of sol-SorCS1 overexpression we never observed any signs of hypoglycemia due to a compensatory decrease in insulin secretion.

**Figure 2:**
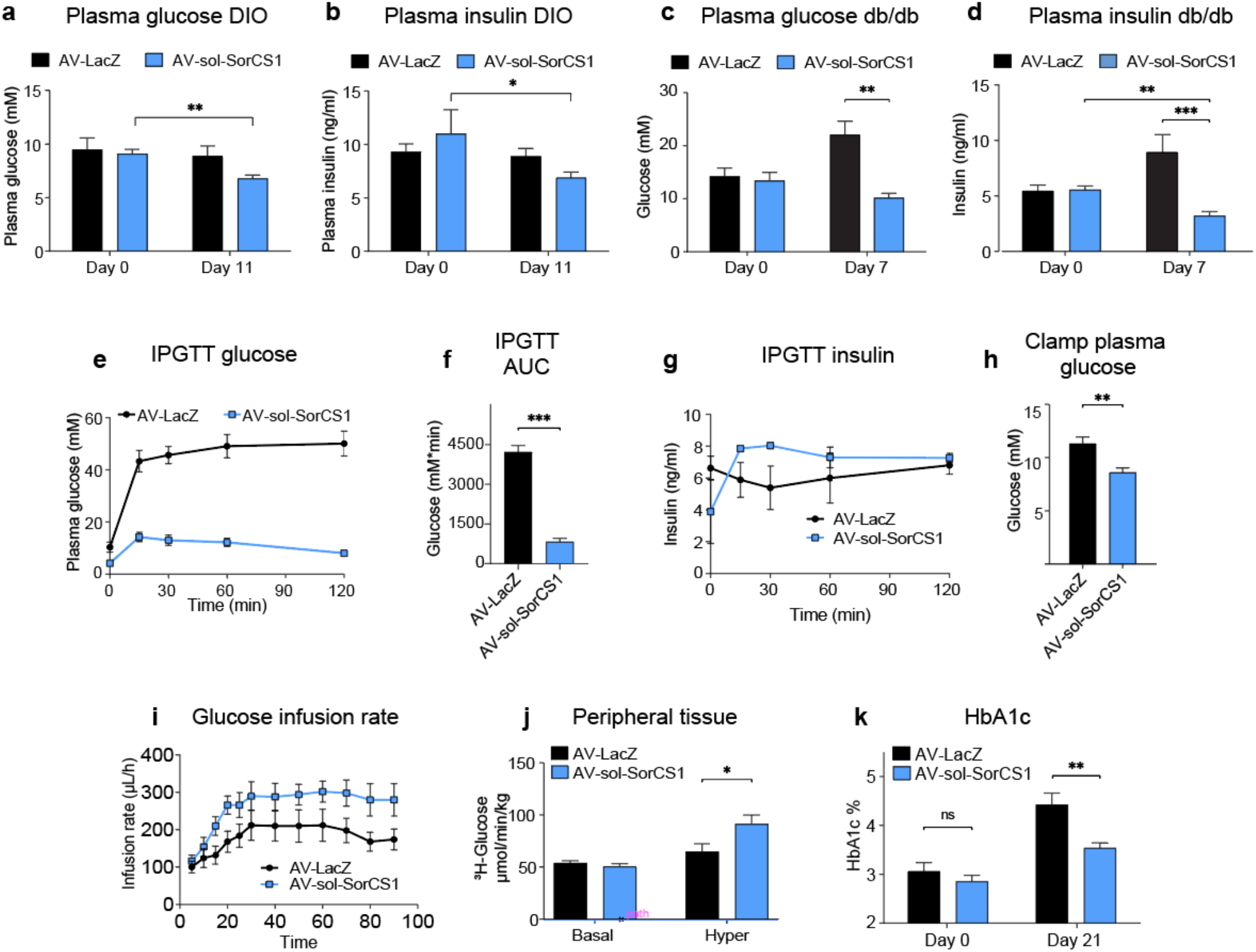
Sol-SorCS1 is an insulin sensitizer. **a**-**d**, treatment with AV-sol-SorCS1, but not AV-LacZ, reduces plasma glucose levels and plasma insulin in 16 weeks old diet-induced-obese (DIO) mice and in 7 weeks old *db/db* mice. **e**, enhanced glucose uptake determined by IPGTT in AV-sol-SorCS1 treated *db/db* mice shown in real-time and, **f**, as AUC (n=4/group). **g**, plasma insulin in the IPGTT shown in e-h. **h**, lower initial fasting plasma glucose in AV-sol-SorCS1 treated *db/db* mice but higher infusion rate during euglycemic hyperinsulinemic clamp, **i** (n=5/group). **j**, the higher infusion rate is accounted for by increased peripheral glucose uptake (primarily muscle tissue). **k**, reduced HbA1c in *db/db* mice 21 days after infection with AV-sol-SorCS1 infection. Data are presented as means ± SEM, * = *P*<0.05, ** = *P*<0.01, *** = *P*<0.001.

Next, we performed hyperinsulinemic euglycemic clamp^19,20^ which is commonly used to assess glucose deposition rates and insulin sensitivity. With the mice clamped to their basal fasting glucose levels (**Fig. 2,h**), the necessary glucose infusion rate to maintain stable plasma glucose levels in AV-sol-SorCS1 mice was substantially higher compared to the AV-LacZ group (**Fig. 2,i**). Notably, sol-SorCS1 increased peripheral glucose uptake by 41%, demonstrating that sol-SorCS1 greatly increases insulin sensitivity in peripheral tissues (**Fig. 2,j**). The difference in severity of diabetes progression was further emphasized when measuring glycated hemoglobin (HbA1c) three weeks after viral injection in another set of mice (**Fig 2,k**). In the AV-LacZ group, HbA1c had increased considerably whereas it was only marginally higher in the AV-sol-SorCS1 group, indicating a long-lasting beneficial effect of the soluble receptor in reducing plasma glucose.

We previously reported that in neurons, the SorCS1 paralogues sortilin and SorCS2, can physically interact through their extracellular domains with the tropomyosin receptor kinase B (TrkB) that binds brain-derived neurotrophic factor (BDNF), to enable its signaling capacity^21,22^. We hypothesized that SorCS1 may likewise engage a tyrosine kinase receptor to promote its activity. Given the genetic association with type 2 diabetes and its glucose lowering function, we asked whether this tyrosine kinase receptor could be the insulin receptor. To assess this, we first performed co-IP experiments between SorCS1 and IR in transfected HEK293 cells in the absence or presence of insulin (**Fig. 3,a**). Antibodies against IR co-precipitated SorCS1 and vice versa, and this interaction was unaffected by preincubation with 100 nM insulin, a concentration that saturates IR. Next, we evaluated the affinity of the receptor-receptor interaction by surface plasmon resonance (SPR) analysis using the extracellular domains of SorCS1, corresponding to sol-SorCS1, and the extracellular domain of IR. We found that sol-SorCS1 bound immobilized IR with an estimated affinity of 5 nM (**Fig. 3,b**). Importantly, the interaction between SorCS1 and IR was not prevented by insulin, as the signals for binding of SorCS1 and insulin to IR separately were additive when the two ligands were co-injected (**Fig. 3,c**). Since insulin did not bind to immobilized SorCS1 (**Extended Data 2)**, the combined data indicate that IR can form a trimeric complex with insulin and soluble SorCS1. Pro-insulin, glucagon, or glucagon-like peptide 1 (GLP-1) also did not bind to immobilized SorCS1 (**Extended Data 2**). We then asked whether full-length SorCS1, when embedded in plasma membrane, may regulate insulin binding to IR and its signaling capacity. First, we overexpressed IR in HEK293 cells (HEK-IR) alone or in combination with SorCS1-b, an isoform of SorCS1 that predominates at the plasma membrane^18^. Remarkably, SorCS1 increased the affinity of insulin to IR by approximately five-fold (**Fig. 3,d**). Given that overexpression of SorCS1 and IR could potentially result in ligand independent autoactivation, we compared insulin signaling in C2C12 cells with or without CRISPR/Cas9 mediated knockout of *Sorcs1*. The most prominent pathway regulating the insertion of GLUT4 containing vesicles is the AKT signaling cascade. Indeed, we found that SorCS1-deficient C2C12 cells were substantially less insulin responsive than control cells as the EC_50_ for pAKT was left-shifted approximately five-fold (**Fig. 3,e and Extended Data 1,b**). Together the data suggested that the glucose lowering effect of SorCS1 is accounted for by an allosteric change in IR that increases its insulin binding affinity, leading to stronger AKT signaling. To substantiate such a model, we measured the thermal stability of full-length IR in the absence or presence of sol-SorCS1 using a cellular thermal shift assay (CETSA)^23^. We found that incubation with sol-SorCS1 right-shifted the melting curve of IR in overexpressing HEK293 cells corresponding to an increase in Tm from 47.8°C to 49.5°C (**Fig 3,f**). The data suggests that sol-SorCS1 induces a conformational change in IR, and supports that allosteric changes in IR may be accountable for the beneficial effects of SorCS1 on insulin binding and signaling.

**Figure 3:**
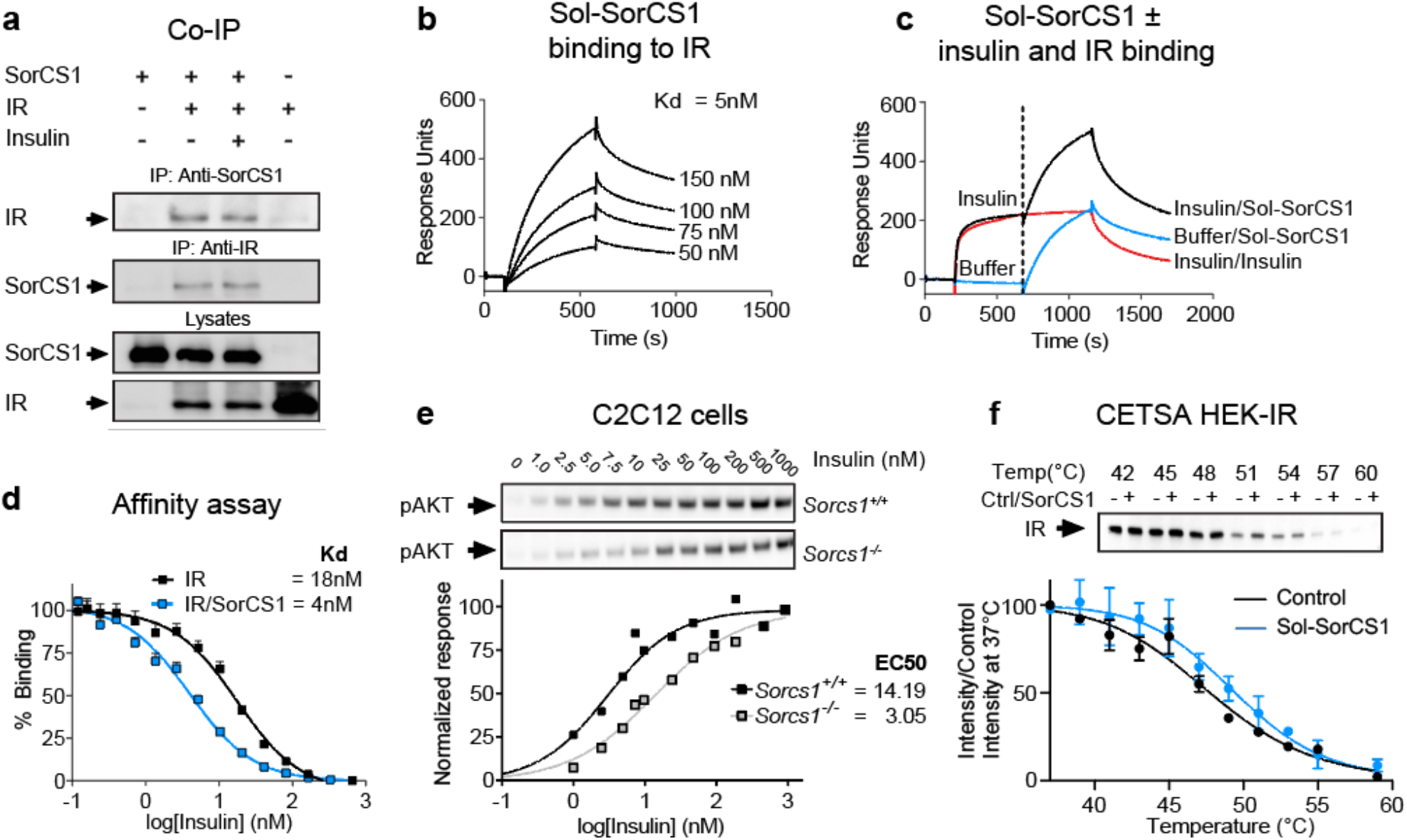
SorCS1 and IR physically interact. **a**, Co-immunoprecipitation between SorCS1 and proIR/IR. The interaction is independent of the presence of saturating concentrations of insulin (100 nM). **b**, SPR analysis demonstrates dose-dependent binding of sol-SorCS1 to IR. **c**, Complex formation of SorCS1, IR, and insulin determined by SPR. At 1200 sec, the additive signals for Sol-SorCS1 (*blue*) and insulin (*red*) equals the response units obtained when IR is incubated with both proteins simultaneously (*black*). **d**, SorCS1 overexpression in HEK293 cells increases insulin affinity to IR. **e**, Reduced phosphorylation of AKT in *Sorcs1*^*-/-*^ C2C12 cells. **f**, 5 min. incubation with 1 µM sol-SorCS1 increases thermal stability of IR from HEK-IR lysate (n=3).

We previously reported that SorCS1 exists as a monomer and as a non-covalent dimer, with the oligomeric state, being regulated by the level of glycosylation^24^. However, the functional implications of the two forms, e.g. whether the relative amounts of the two forms dictate binding specificity remains unknown. To explore the spatial distribution of the monomer and dimer at the subcellular level, we used a bimolecular fluorescence complementation (BiFC) assay^25^. HEK-IR cells were transfected with two human SorCS1 constructs each containing one-half of the Venus fluorescent protein. When SorCS1 dimerizes the two Venus fragments unite to elicit a green fluorescent signal that echoes the subcellular distribution of dimeric SorCS1. First, we stained the cells with an antibody that recognizes both monomeric and dimeric SorCS1 and found monomeric SorCS1 is present at the plasma membrane whereas the BiFC-positive signal representing dimeric SorCS1 is restricted to intracellular compartments (**Fig. 4,a**). This is in accordance with our previous observation demonstrating that through its maturation, N-linked oligosaccharides transforms SorCS1 from the dimeric form into the monomeric configuration^24^. Binding of insulin to the IR changes its dimeric and symmetric conformation to adapt an asymmetric configuration (personal communication, Dr Poul Nissen, manuscript to be resubmitted). This prompted us to study whether the interaction of IR is restricted to the monomeric form of SorCS1. To demonstrate that monomeric SorCS1 and IR can interact, we first combined the BiFC signal with proximity ligation assay (PLA) for the two receptors. In this assay a red fluorescent signal appears when SorCS1 and IR are in close proximity (<40 nm). We observed a strong PLA signal at the plasma membrane that was BiFC-negative, indicating that only monomeric SorCS1 binds IR, which suggest that the biologically active form of SorCS1 is the monomer (**Fig. 4,b)**. To substantiate this hypothesis, we produced an adenovirus encoding a truncated version of SorCS1 that is unable to dimerize and studied its biological activity. SorCS1 harbors a consensus sequence (amino acids 695 to 702) for cleavage by prohormone convertases of the subtilisin/Kex2-like family, such as furin^26^. Truncation at this site gives rise to a soluble fragment of roughly 60 kDa (aa130-695 after cleavage of the propeptide and signal peptide) that comprises the entire Vps10p domain but does not possess the dimerization site (**Extended Data 3)**. We overexpressed this explicit monomer by adenovirus (AV-aa1-695-SorCS1) and compared its glucose lowering capacity to AV-sol-SorCS1 (aa1-1097) that comprises the entire ectodomain and likely represents a mixture of monomer and dimer. At day 7 and 14 after viral injection both soluble SorCS1 variants were equally potent in reducing plasma glucose (**Fig. 4,c**). This finding indicates that shed monomeric sol-SorCS1 plays an important role in the biological effects on peripheral glucose disposal.

**Figure 4:**
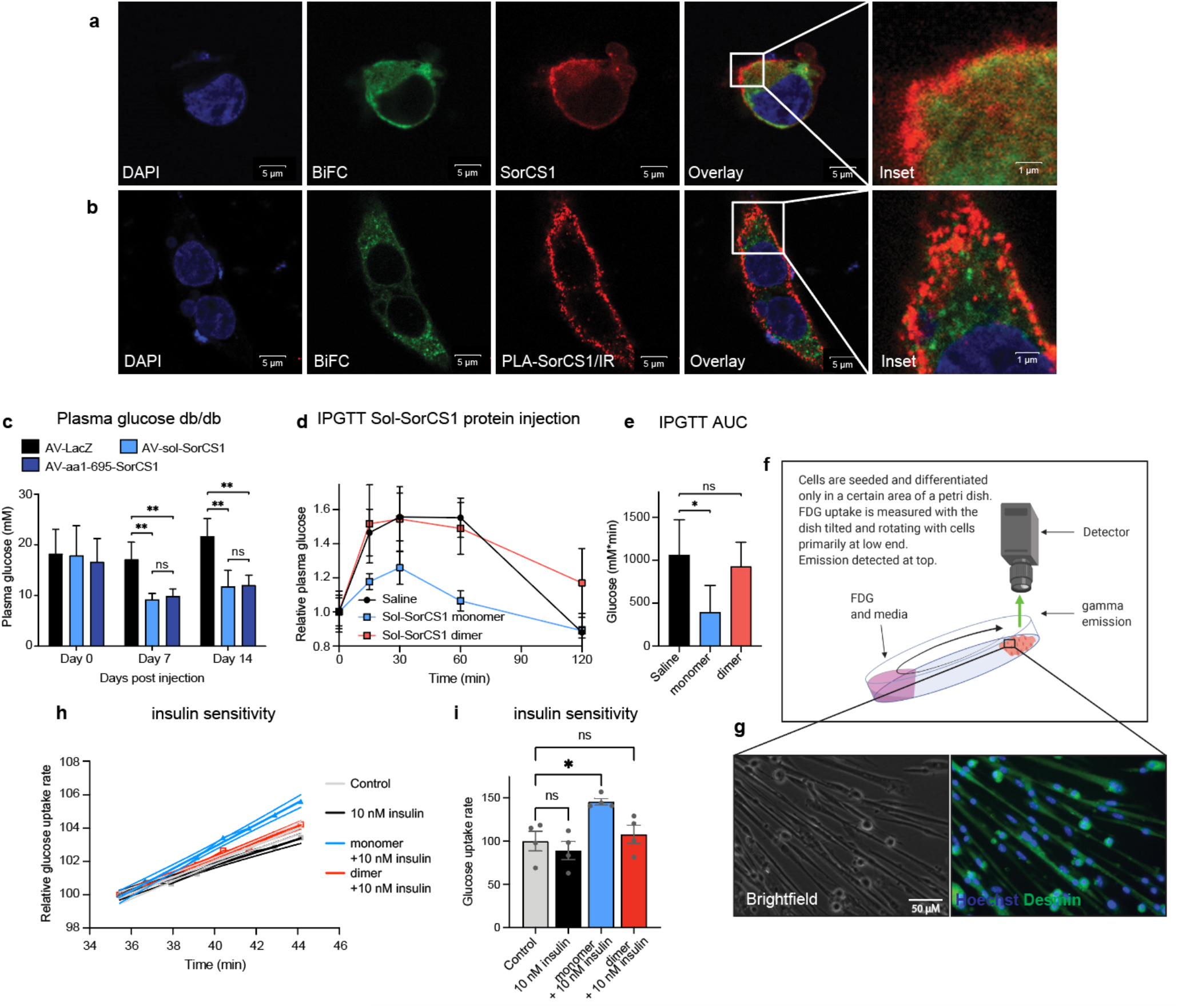
**A**, SorCS1 dimers that are BiFC-positive (*green*) are confined to cytosolic compartments, whereas monomeric and BiFC-negative SorCS1 (*red*), is present at the cell surface (**a**, *inset*). **b**, in non-permeabilized HEK293, PLA for SorCS1 and IR (*red*) demonstrates binding between monomeric SorCS1 (BiFC-negative) and IR. **c**, AV-sol-SorCS1 and AV-aa1-695-SorCS1, but not AV-LacZ, reduces plasma glucose in 8 weeks old *db/db* mice. **d**, intraperitoneal injection of recombinant monomeric, but not dimeric, sol-SorCS1 increases glucose clearance in *db/db* mice. **e**, AUC of the data displayed in panel **d. f**, illustration of kinetic FDG uptake assay. **g**, representative brightfield and fluorescence image of the murine myotubes used for the uptake study. Hoechst in *blue* and Desmin in *green*. **h**, relative FDG uptake in control cells (gray), 10 nM insulin (black), sol-SorCS1 monomer 10 + nM insulin (blue) and sol-SorCS1 dimer 10 + nM insulin (red) (n=4). **i**, bar chart overview of the slopes displayed in panel **h**. statistics are done on actual slopes. Data are presented as means ± SEM, * = *P*<0.05, ** = *P*<0.01.

To demonstrate that the beneficial properties of sol-SorCS1 was not secondary to intrahepatic effects upon hepatic overexpression and to clearly isolate the actions of monomeric from dimeric sol-SorCS1, we studied the glucose lowering effect of the two species upon exogenous administration. In practice, the two forms were purified by size exclusion chromatography from medium of cells expressing sol-SorCS1 and the two species injected intraperitoneally into *db/db* mice at 10 hrs and 1 hr prior to a glucose tolerance test. Intriguingly, monomeric sol-SorCS1 profoundly improved glucose tolerance whereas the dimeric version showed no effect on glucose disposal (**Fig. 4,d**). The AUC for plasma glucose in the glucose tolerance test was reduced by approximately 60% (from 1060 to 393 mM*min, P<0.05) in animals treated with monomeric sol-SorCS1 (**Fig. 4,e**).

Aimed at demonstrating a direct effect of monomeric sol-SorCS1 on muscle cells, we developed a real time *in vitro* assay with a 1 minute resolution in cellular glucose-uptake (**Fig. 4,f-g, Extended Data 4**)^27^. Applying this highly sensitive real-time assay on primary murine myotubes, revealed that sol-SorCS1 does not function as an insulin mimetic, since preincubation with either variant of sol-SorCS1 for 20 minutes did not increase uptake of the glucose analogue (**Extended Data 4,b**). Subjecting naïve myotubes to 10 nM insulin had no effect on glucose analogue uptake. However, after preincubation with 100 nM monomeric sol-SorCS1 addition of 10 nM insulin led to a 45% increase in glucose uptake rate (**Fig 4,h-i**). In marked contrast, 10 nM insulin stimulation after preincubation with dimeric sol-SorCS1 which showed no such effect. However, addition of 100 nM insulin to naïve or dimeric sol-SorCS1 treated myotubes increased glucose uptake to match the rate obtained for myotubes stimulated with 10 nM insulin and pretreated with the sol-SorCS1 monomer (**Extended data 4,c)**. Hence, monomeric sol-SorCS1 acts directly on myotubes to enhance their insulin sensitivity and induces a rapid increase in glucose uptake.

In conclusion, we have demonstrated that SorCS1 circulates in plasma in a soluble form that can be produced by adipose tissue. Once secreted it acts as a hormone that targets tissues remote from its site of production to increase insulin sensitivity. Accordingly, overexpression of the soluble receptor fragment reduces plasma glucose and insulin levels in mice with diet induced obesity and limits diabetes progression in leptin receptor knockout mice. We provide evidence that the biological activity of sol-SorCS1 is restricted to its monomeric form and suggest that once liberated from the plasma membrane it acts in an endocrine fashion (**Extended Data Fig. 5)**. Our study nominates sol-SorCS1 as a novel adipokine to control peripheral glucose metabolism and insulin sensitivity, and as a template for future anti-diabetic drug development.

## Acknowledgements

The study was funded by the Lundbeck Foundation grant no. R248-2017-431 (AN), DANDRITE-R248-2016-2518 (AN), R90-2011-7723 (AN); The Independent Research Fund Denmark grant no. DFF-7016-00261 (AN); The Novo Nordisk Foundation grant no. NNF16OC0020984 (AN); The Danish National Research Foundation grant no. DNRF133 (AN); The Danish Diabetes Academy, Novo Nordisk Foundation (MK), Lundbeck Foundation Emerge (MFK, AN). We thank Benedicte Vestergaard, Anja Aagaard and Anne Kerstine Thomassen for excellent technical assistance and Professor Alan Attie, University Wisconsin-Madison, WI, for insightful discussion.

---

**Extended Data Figure 1:**
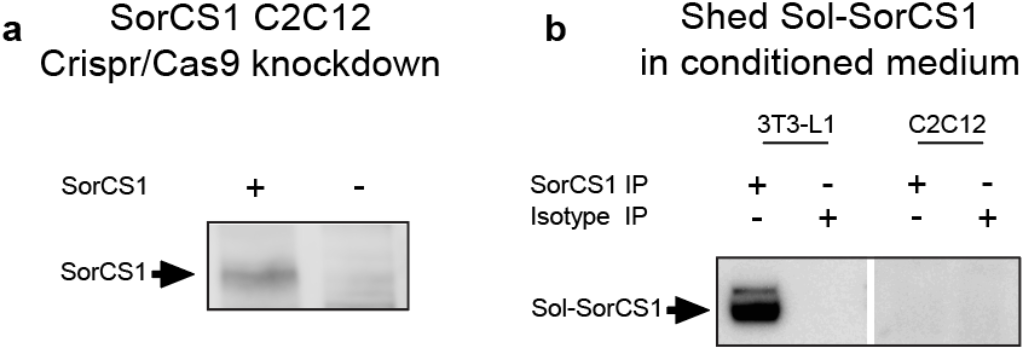
**a**, Through Crisp-Cas9 gene editing, we created a stable SorCS1 knockdown model of the murine C2C12 muscle cell line. SorCS1 protein presence was validated in the wild-type cells by western blotting using a single homemade polyclonal antibody targeting murine SorCS1 (MK-S1). **b**, SorCS1 could not be detected in serum free conditioned medium (hybridoma) from C2C12 cells using two polyclonal antibodies, whereas it was clearly present in conditioned medium from 3T3-L1.

**Extended Data Figure 2:**
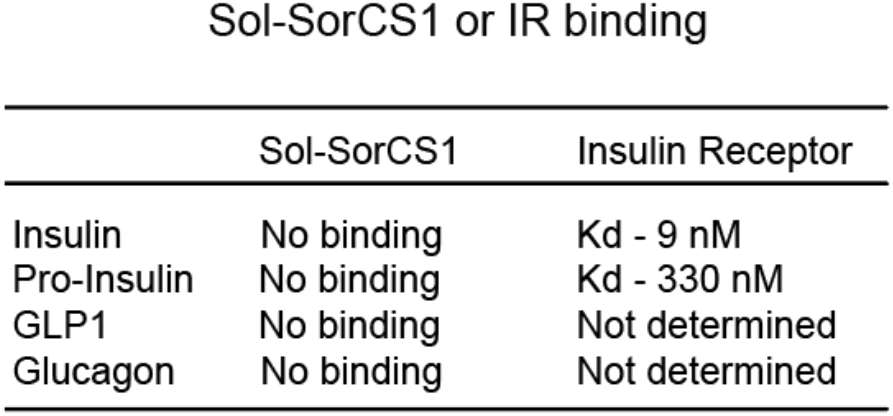
Table showing no binding between sol-SorCS1 and various key factors in peripheral plasma glucose clearance.

**Extended Data Figure 3:**
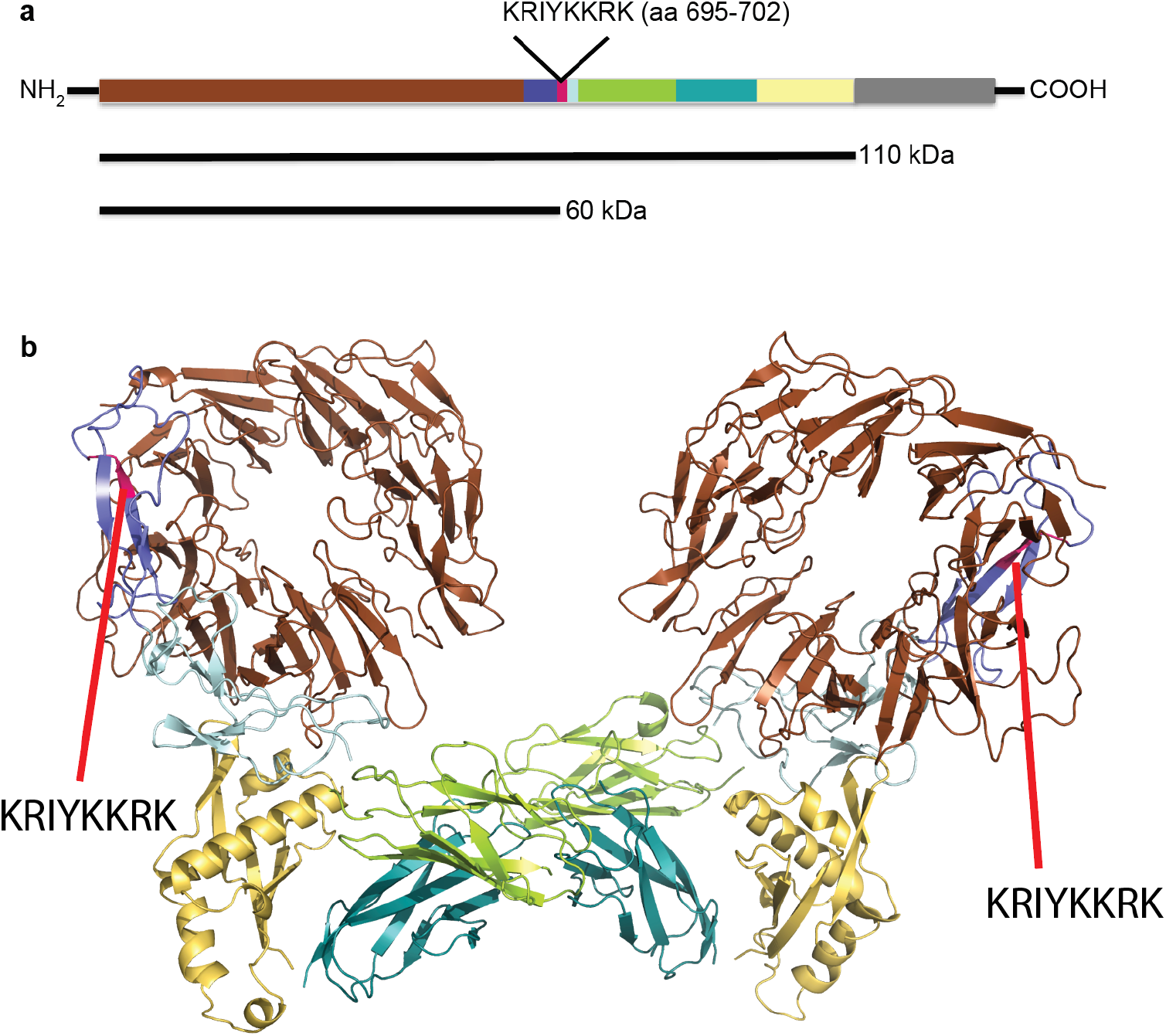
**a**, illustration of potential furin site on SorCS1 in dimer formation and the termination site of the explicit monomer “aa1-695-sol-SorCS1”. **b**, A homology model of human dimeric SorCS1 based on murine SorCS2 (PDB entry 6FG9)^28^ constructed using the SwissModel service^29^. The individual domains were manually overlaid on the single particle reconstructed envelope of monomeric SorCS1 (EMDB entry EMD-3708)^24^. The SorCS1 dimer: Colors: β-propeller – *brown*, 10CCa – *blue*, 10CCb – *lightblue*, PKD1 – *green*, PKD2 – *cyan*, and SoMP – *yellow*. The position of the putative furin site (699-KKRK-702) delimiting the sol 1-702-SorCS1 construct is indicated in pink.

**Extended Data Figure 4:**
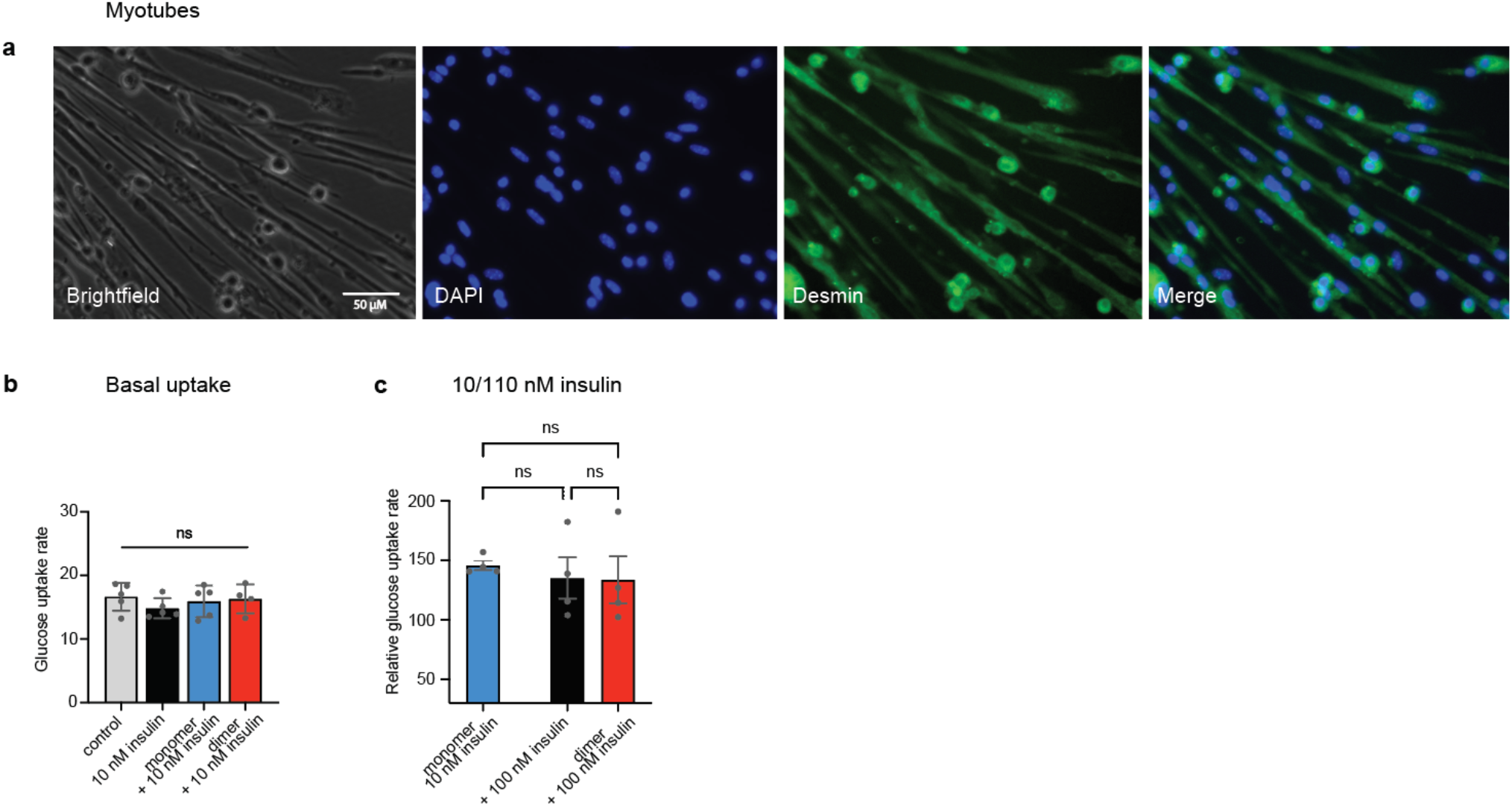
**a**, All individual panels from brightfield and fluorescence image Fig 4,h. **b**, Basal glucose analogue uptake rate showing no difference between interventions on each study day. **c**, naïve myotubes and myotubes pretreated with 100 nM dimeric sol-SorCS1 increase glucose analogue uptake only when stimulated with an additional 100 nM insulin compared to 10 nM insulin in myotubes pretreated with 100 nM monomeric sol-SorCS1.

**Extended Data Figure 5:**
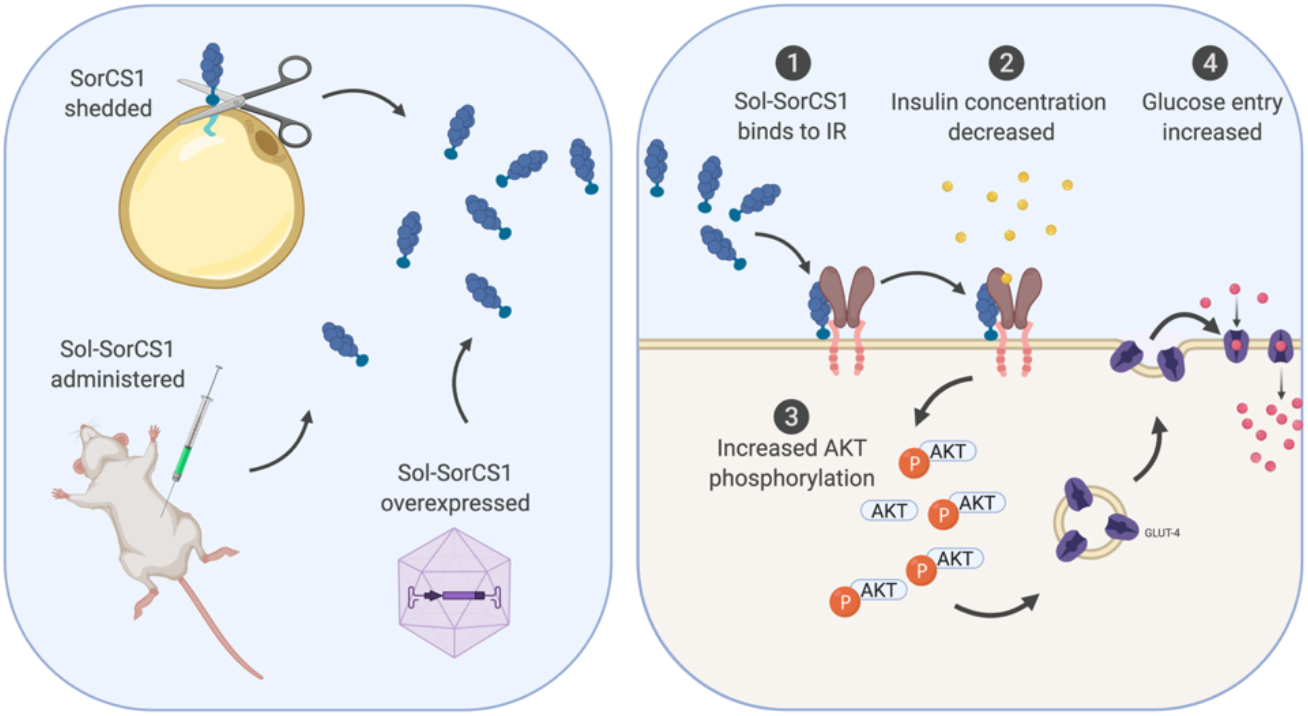
An illustrated summery of mode of action of sol-SorCS1. The illustration shows how shedding, injection or overexpression of sol-SorCS1 leads to increased glucose uptake through increased insulin receptor signaling while circulating insulin concentrations decrease implying increased receptor sensitivity.

