## Supplementary material for "Soluble SorCS1 binds the insulin receptor to enhance insulin sensitivity": Methods

[1]PROMEMO and DANDRITE, Aarhus University, Denmark; [2]Department of Clinical Pharmacology, Aarhus University Hospital, Denmark and Danish Diabetes Academy; [3]Department of Biomedicine, Aarhus University, Denmark; [4]Department of Molecular Biology, Aarhus University; [5]Novo Nordisk Foundation Center for Basic Metabolic Research, Copenhagen University, Denmark; [6]Department of Neuroscience, Mayo Clinic, Florida, USA; 7 Hamburg Center of Neuroscience (HCNS), Hamburg, Germany; [8]Osnabrück University, Frankfurt Am Main Area, Germany; [9]Lund University Diabetes Centre, Malmö, Sweden; [10]Dept. Medicine, Div. Endocrinology, and Einthoven Laboratory for Experimental Vascular Medicine, Leiden University Medical Center (LUMC), Leiden, The Netherlands; [11]RMPC, Department of Biomedical Sciences, Copenhagen University, Denmark;

* Shared first authors. #Corresponding authors

### Human tissue and cohort description

Human pancreatic islets (n = 188), liver (n = 12), fat (n = 12), and skeletal muscle (n = 12) of European Ancestry were obtained from organ donors through the EXODIAB network from the Nordic Islet Transplantation Program (http://www.nordicislets.org). All procedures were approved by the ethics committee at Lund University and performed as described[30]. Total RNA was isolated with the AllPrep DNA/RNA Mini Kit (Qiagen). RNA quality and concentration were measured using an Agilent 2100 bioanalyzer (Bio-Rad) and a Nanodrop ND-1000 (NanoDrop Technologies), respectively. Raw data were base-called and de-multiplexed using CASAVA 1.8.2 (Illumina) before alignment to hg19 with STAR. Gene and exon feature count of the number of reads was performed using featureCounts (version 1.4.4, http://bioinf.wehi.edu.au/featureCounts/). Counts were normalized for library sizes using the trimmed mean of M-values (TMM) and transformed into log2 counts per million (log2CPM) using *voom* (limma R, Bioconductor) implemented in edgeR. Combat was used for removal of batch effects using the Combat function (sva package). A linear model adjusting for age, sex and purity was used to determine the expression of genes in relation to quantitative phenotypes like BMI[31].

Plasma concentration of sol-SorCS1 was measured in 481 individuals of whom 120 had diabetes. Three individuals had fasting plasma glucose levels below 2.2 and these individuals (3 diabetics) were removed from analyses of fasting plasma glucose and traits including fasting plasma glucose.

Human adipose explant tissue was acquired through liposuction from anonymous healthy individuals. The tissue was thoroughly rinsed in saline, pre-incubated overnight prior to 24 h incubation at 37°C in fresh media consisting of: M199 (Sigma), 1% BSA (calbiochem), P/S (Gibco), hepes (Sigma) 1.1 g $NaHCO_3$ (Merck), Glutamax (Gibco), 1 nM insulin (Sigma) 2 µg/ml Antipain (Sigma) Leupeptine (Sigma), as previously described[32]. The incubation was stopped by carefully collecting the media avoiding the floating adipose tissue. A low-speed centrifugation was used to separate any remaining floating fat cells and pelleted non-adipose cells and the media was collected, cooled, sterile filtered and concentrated by a 50 kDa cut-off Amicon Ultra-15 Centrifugal Filter Units (Merck).

**Custom made ELISA against human soluble SorCS1**

96-well plate Nunc MaxiSorp were coated with anti-hSorCS1 (Dako F7050, custom made against EC-Domain) 1 µg per well in 55 mM $NaHCO_3$ pH 9.8. The wells were subsequently blocked with 5% BSA (Sigma A7888) in PBS (pH 7.4). Washing was done in PBS-T (pH 7.4). Sample, internal control and standard was incubated overnight at 4°C. Standard curve was made from 0.625 to 160 ng/ml of hSorCS1 (AF3457SR, R&D Biosystems). The detection antibody (AF3457 from R&D, Goat-anti-hSorCS1) was diluted in PBS-T with 1% BSA to 1 µg/ml. 100 µl of the solution was added to each well, and allowed to incubate at 37°C for one hour. The secondary antibody (Dako P0160 Rabbit-anti-Goat-HRP) was diluted in PBS-T with 1% BSA, 1:1000. 100 µl of the solution was added to each well followed by incubation at room temperature for one hour. The plate was developed by addition of OPD (Dako S2045) in $ddH_2O$. The reaction was stopped with 2 M sulphuric acid. Absorbance was read at 490 nm wavelength.

**Western blotting**

Protein concentration was measured using the Bio-Rad Protein Assay extraction method. Lysates were mixed with SDS sample buffer boiled at 95°C for 5 min and subjected to reducing SDS-PAGE using, 4-12%, 8%, or 4-16% acrylamide gels as previously described[33] unless stated otherwise.

**Antibodies**

The purified soluble extracellular domain of human SorCS1 (SorCS1 ECD) was used to generate custom-made rabbit polyclonal anti-SorCS1 antibodies (αECD) (Dako F7050), and chicken polyclonal anti-SorCS1 aECD (MK-S1) as well as polyclonal antibodies against leucin-rich domain of murine SorCS1. Other primary antibodies used were anti-SorCS1 (AF3457, R&D), rabbit anti-IR (sc-711, Santa Cruz or 07-724 Millipore/Sigma), anti-Akt (9272, Cell Signaling), anti-pAkt (Ser473) (9271s, Cell signaling), anti-Desmin (D1033, Sigma), mouse anti-β-actin (AF5441, Sigma), and anti-GAPDH (G8795, Sigma).

**Co-immunoprecipitations of soluble SorCS1 and IR**.

Co-immunoprecipitation studies on HEK293 cells overexpressing the human insulin receptor and SorCS1 using DSP crosslinker. Antibodies used for immunoprecipitation studies were anti-IR sc711, C-19 (Santa Cruz Biotechnologies), 07-724 (Millipore/Sigma), anti-SorCS1 AF3457 (R&D) and a custom-made antibody against the ECD of hSorCS1. The immunoprecipitation method has been described previously[33]. Precipitation was done using protein G Sepharose (Roche Applied Science).

**Animal studies**

Leptin receptor deficient mice (db/db) mice (BKS.Cg/BomTac-m+/+Lepr<db>), diet induced obese mice (DIO-B6) and wild-type controls (C57BL/6-BomTac) were procured from Taconic (Denmark).
All mice were housed in an environmentally controlled facility (12-hour light and dark cycles) with free access to food (1324; Altromin (unless otherwise stated)) and water.

**Blood glucose and plasma insulin measurements.**

Mice were fasted overnight from 10 PM to 8 AM. Blood samples were obtained by retro-orbital bleeding after induction of anesthesia with isoflurane (IsoFloVet, Orion Pharma, Denmark). Plasma glucose was measured (Ascensia Contour, Bayer, Germany). The blood samples were collected in 0.5 ml lithium-heparin tubes (Sarstedt, Germany). The tubes were centrifuged for 15 min at 3,000xg, and the plasma was separated and stored at -20°C until assayed. Plasma insulin levels were determined using a mouse insulin ELISA (DRG Diagnostics, Marburg, Germany). Blood samples from conscious mice were obtained through cheek and tail blood sampling. HbA1c was measured in full blood (Cat# 80310, Crystal Chem, The Netherlands).

**Glucose tolerance test.**

Intraperitoneal glucose tolerance tests (IPGTT) were performed using fasted, age matched mice. Each mouse received an intraperitoneal (IP) bolus of D-glucose (Sigma) in sterile saline. Blood samples, <30 µl, were obtained at times 0, 15, 30, 60, and 120 min after injection.

**Hyperinsulinemic isoglycemic clamp procedure in db/db mice.**

Basal and insulin-stimulated glucose metabolism was determined in postabsorptive (i.e., overnight fasted 10 PM to 9 AM), body weight–matched mice[34]. Animals were anesthetized with 6.25 mg/kg acepromazine (Plegicil, Dechra, Denmark), 6.25 mg/kg midazolam (Roche, Denmark), and 0.31 mg/kg fentanyl (Janssen-Cilag, Denmark). Tissue-specific glucose uptake was determined in hyperinsulinemic isoglycemic state. In the basal period the db/db mice either transfected with AV-LacZ or AV-Sol-Sorcs1 7 days earlier, were infused with [$^3$H]glucose (0.3 µCi/kg/min) for 60 min to attain steady state levels of [$^3$H]glucose in plasma, and to determine endogenous glucose production and clearance. In the hyperinsulinemic isoglycemic clamp period, insulin was administered intravenously by a primed dose (4.1 mU), followed by continuous (20 mU/kg/min)[35] infusion to attain steady-state insulin levels, together with [$^3$H]glucose (0.3 µCi/kg/min) for 90 min[36]. Intravenous infusion of a 12.5% d-glucose solution was used to maintain isoglycemia as determined at 5 min intervals the first 30 min of the hyperinsulinemic state, hereafter in 10-min intervals, all via tail bleeding (<3 µl, Contour NEXT, Bayer Group, Germany).

### Cell Studies

#### Isolation and differentiation of myotubes from murine muscle.

Primary murine satellite cells (MuSC) were sorted by Fluorescence Activated Cell Sorting (FACS)[37], cultured and differentiated from the gastrocnemius of 9 weeks old wild-type C57BL6/BomTac mice. In short, muscles were dissected and minced in phosphate-buffered saline (PBS) (Invitrogen), dissociated mechanically with sterile scissors and enzymatically in a solution of dispase (grade II, 2.4 U/ml; Roche Molecular Biochemicals) and collagenase (class D, 1%; Roche Molecular Biochemicals). The slurry was maintained at 37°C with intermittent mixing. The enzymatic solution was removed by sedimenting the dissociated cells by centrifugation. The pellet was then resuspended in growth medium filtered through a 70 µm and a 30 µm mesh before finally being subjected to FACS. MuSC was sorted by negative selection on CD31, CD45, Ter-119 and Sca1 and positive selection on integrin a7 (all Miltenyi Biotec)[37]. After sorting, cells were plated and cultured in a humidified incubator at 37°C in 5% $CO_2$, in a 9.6 $cm^2$ dish precoated with diluted (1:1000) Engelbreth-Holm-Swarm murine sarcoma (ECM) gel (Sigma-Aldrich) in proliferation media (High glucose DMEM (Lonza), 20% fetal bovine serum (Sigma Aldrich), penicillin-streptomycin (Gibco)) supplemented with fresh fibroblast growth factor basic (bFGF) (Thermo Fisher Scientific (2.5 ng/ml). MuSCs were split at 80% confluency and replated on ECM coated dishes. MuSCs were differentiated into myotubes at 90-100% confluency by omitting bFGF and reducing FBS to 2%. Differentiation was confirmed by visual inspection before assessing glucose uptake.

**Preparation of Adenoviruses and soluble protein**

The recombinant adenovirus for expression of human soluble SorCS1 (hsol-SorCS1, aa1-1097 and sol-SorCS1, aa1-695) was generated as follows: pcDNA3.1/Zeo(−)/hsol-SorCS1 encoding the human soluble *SORCS1* cDNA (amino acids 1-1100) was digested with Pme1 and Apa1 and the fragment encoding hsol-SorCS1 inserted into the shuttle plasmid pVQpacAd5CMVK-NpA (ViraQuest, North Liberty, IA) generating the plasmid pVQAd5CMVK-NpA-sol-SorCS1. The plasmid was sent to ViraQuest for generation and propagation of adenovirus expressing hsol-SorCS1. A negative control virus, VQAd5 CMV ntLacZ, that expresses beta-galactosidase, was purchased from ViraQuest. Animals were injected with 2E9 PFU per mouse[33] in 100 µl sterile isotonic saline in the tail vein. Human and murine SorCS1 protein was produced by stably transfected Chinese hamster ovary (CHO-K1) cells. The monomeric and dimeric soluble SorCS1 proteins were purified by Superdex 200 size exclusion chromatography and found >90% pure as previously described[38]. For *in vivo* injections, 80-100 µg protein was injected IP at each time point. For *in vitro* incubations, 100 nM concentrations were used.

**Plasmids**

Human SorCS1a, SorCS1b, SorCS1c were used, and have previously been described[39]. Bimolecular fluorescence complementation SorCS1 plasmids were generated with ECD and TM domain of SorCS1, a tag (HA or V5), a linker and 1 half of the Venus fluorescence molecule[40]. The Venus molecule is separated into two parts that are not fluorescent by themselves. N-Venus is aa1-158, and C-Venus is aa159-239. When the two parts (N- and C-) are combined, it yields a fluorescent molecule (Ex515/Em528). The two constructs named hSorCS1-N-Venus (1-158) and hSorCS1-C-Venus (159-239) can exist as monomers or dimers. The dimers (N and C, not N-N or C-C) will form a fluorescent complex[40].

**Cell lines**

C2C12 cells (ATCC, CRL-1772) were differentiated in DMEM with 5% heat inactivated horse serum. *mSorCS1* was targeted in C2C12 cells using Origene CRISPR/Cas9 murine SorCS1 kit (KN316472) according to manufacturer's protocol, with Sorcs1 gRNA vector 1 in pCas-Guide vector (KN316472G1), Sorcs1 gRNA vector 2 in pCas-Guide vector (KN316472G2), and donor vector (KN316472D) containing left and right homologous arms and GFP-Puro functional cassette. Control cells were passaged alongside using scrambled gRNA vector (SKU GE1000003). The cells were tested in western blot analysis to ensure complete knockout. HEK293 (ATCC CRL-1573) were stably transfected with human IR (long) including exon17 (HEK-IR), and/or human SorCS1.

**Surface plasmon resonance (SPR) analysis**

SorCS1 and IR interaction was interrogated using surface plasmon resonance analysis in a BiaCore system. Soluble murine SorCS1 (custom made or R&D 3457-SR-050) and insulin receptor (R&D 1544-IR-050/CF) were immobilized on a CM5 chip. Sample and running buffer were 10 mM HEPES, 150 mM $(NH_4)_2SO_4$, 1.5 mM $CaCl_2$, 1 mM EGTA, 0.005% Tween-20 (pH 7.4). The SPR signal was expressed in relative response units (RUs) after subtraction of the RU in a control flow channel (reference flow cell). Kinetic parameters were determined using BIAevaluation 3.1 software. Saturating concentrations of insulin (500 nM) was injected to the IR-coupled biosensor chip at t=200 sec and t=580 sec (Purple curve on Fig 3c). After 580 sec 100 nM sol-SorCS1 was added to the incubation buffer and binding to IR was monitored (blue curve on Fig 3c). Black curve in Fig 3c: Incubation with insulin at t=200 sec, followed by co-injection of insulin and SorCS1 at 580 sec. After 1,200 sec the combined signals from insulin (red) and SorCS1 (blue) separately, corresponds to the signal obtained by simultaneous incubation with insulin and SorCS1 (black).

**Insulin competition binding in cell models.**

HEK293 cells stably overexpressing either insulin receptor or both insulin receptor and SorCS1, were seeded in poly-lysine coated 24-well trays, and incubated for 48 h. Cells were then cooled on ice, before adding 80 pM $^{125}$I-labelled insulin (Perkin Elmer, NEX420010UC) and non-labeled insulin in the range of 40 pM – 320 nM. The cells were then incubated at 5°C in a HEPES buffer (124 nM NaCl, 4.7 mM KCl, 2.5 mM $CaCl_2$, 1.2 mM $MgSO_4$, 25 mM HEPES, 0.2% BSA; pH 7.4). The buffer was collected after 6 h and cells were dissolved in 0.5 M NaOH and collected in a separate tube. All tubes were counted for gamma radiation to produce competition binding curves. Log transformed insulin concentrations and binding percent were fitted with sigmoidal dose response nonlinear fit, using GraphPad Prism 9 software.

**Cellular Thermal Shift Assay (CETSA)**

Lysates from HEK-IR cells were washed x2 in PBS and harvested in a 100 mM HEPES (pH 7.5) + 1 mM DTT + 1x "Complete protease inhibitor cocktail" and subjected to x4 freeze thaw cycles using dry ice and centrifuged at 20,000 x g for 20 minutes at 4°C. CETSA was performed in accordance with the Molina *et al*. but with sol-SorCS1 replacing the small molecule. In short, lysate was diluted to 1 µg/µl and 350 µl was added to two separate tubes[41]. Next, 20 mM $MgCl_2$ was added alongside 1000 nM sol-SorCS1 (AF3457SR, R&D Biosystems) or PBS as control, or 1000 nM purified sol-SorCS1 monomer or $H_2O$ as control and the tubes were left for 5 minutes at room temperature. The samples were then divided into 30 µl matched aliquots and subjected to 3 minutes heat treatment at individual temperatures followed by 3 minutes cooling at room temperature. Lysates were then centrifuged at 20,000 x g for 20 minutes to sediment the denatured proteins before 10 µl supernatant was transferred in pairs to an SDS-PAGE gel for electrophoresis followed by a western blot for hIR (Cell Signaling, CS3025).

**Immunofluorescence Analysis**

Cells were fixed in 4% paraformaldehyde, permeabilized with 0.1% Triton X-100, blocked in 10% normal donkey serum (TriChem, Jackson Immuno Research), and incubated with primary antibodies against the specific target for 24 h at 4°C. Secondary fluorescent-labeled antibodies were used for visualization (Alexa Fluor, Thermo Fisher Scientific). Specific antibodies used were: anti-insulin receptor (sc711, Santa Cruz, or af1478, R&D), anti hSorCS1 (AF3457, R&D), and anti-desmin (D1033, Sigma Aldrich).

**Proximity ligation assay**

Proximity ligation assay (PLA) (Duolink®II, Olink Bioscience) was performed in accordance with the manufacturer's protocol. Primary antibodies were anti-SorCS1 (R&D Systems AF3457) and anti-insulin receptor β (Santa Cruz Biotech, sc-711). Corresponding oligonucleotide-conjugated secondary antibodies were used for ligation and amplification and nuclei were visualized with DAPI staining. HEK293 cells stably transfected with insulin receptor (HEK-IR), and transiently transfected with SorCS1 were seeded on coverslips and incubated 24-38 hours before fixation in 4% paraformaldehyde for 20 min. For staining of non-permeabilized cells, incubation with SorCS1 primary antibody (R&D Systems AF3457) for 2h at 4°C was followed by fixation in 4% paraformaldehyde[33].

***In vitro* kinetic glucose uptake studies**

To assess insulin-stimulated glucose uptake +/– sol-SorCS1 we developed a new kinetic assay thoroughly described in Breining et al.[42]. In short, murine myotubes were assessed ±insulin (Sigma, #I5500) at two concentrations (10 nM and 110 nM), with or without 20 minutes of preincubation with sol-SorCS1 monomer or dimer. Uptake of the $^{18}$Fluor analogue of 2-deoxy-glucose (2-DG); $^{18}$Fluoro-2-deoxy-glucose (FDG), was used as a proxy for glucose uptake. For each plate included (n=4 and 4 plates per setup), basal FDG uptake was assessed prior to injection of 3 µl containing either 10 nM insulin in LG-DMEM media, or media control, and subsequently an additional 3 µl containing 100 nM insulin in LG-DMEM media, or media control. Uptake was assessed as emission from the myotubes and analyzed[42]. Myotubes were stained in the petri dish and imaged via an inverted microscope (EVOS M5000, Invitrogen)

**Ethics**

All experiments were approved by the Danish Animal Experiments Inspectorate under the Ministry of Justice (Permit 2011/561-119 and 2006/561-1206) and carried out according to institutional and national guidelines. Ethical approvement for the collection of human pancreatic islets, muscle, fat and liver samples from cadaver donors has been obtained from the ethical committee in Lund (Dnr. LU 2011/263)[31]. Human adipose explant tissue was collected fully anonymously (approved by the Central Denmark Region ethics committee).

**Statistics**

Data are presented as means ± SEM, *P<0.05, **P<0.01, ***P<0.001. AUC and peak were calculated on individual curves (IPGTT). Unless mentioned otherwise, significance was evaluated using Students' two-tailed t test, and two-way ANOVA in time-curves. Statistical analysis in GIR was done under steady state conditions. Log transformation was done if needed to achieve Gaussian distribution. A non-parametric approach (Friedman test) with Dunn's multiple comparisons test was used for glucose uptake analysis. Calculations were done using GraphPad Prism software 9.
